# Developmental Morphogens Direct Human Induced Pluripotent Stem Cells Towards an Annulus Fibrosus-Like Cell Phenotype

**DOI:** 10.1101/2022.05.06.490483

**Authors:** Ana P. Peredo, Tonia K. Tsinman, Edward D. Bonnevie, Xi Jiang, Harvey E. Smith, Sarah E. Gullbrand, Nathaniel A. Dyment, Robert L. Mauck

## Abstract

Therapeutic interventions for intervertebral disc herniation remain scarce due to the inability of endogenous annulus fibrosus (AF) cells to respond to injury and drive tissue regeneration. Unlike other orthopaedic tissues, such as cartilage, delivery of exogenous cells to the site of annular injury remains underdeveloped, largely due to a lack of an ideal cell source and the invasive nature of cell isolation. Human induced pluripotent stem cells (iPSCs) can be differentiated to specific cell fates using biochemical factors and are, therefore, an invaluable tool for cell therapy approaches. While differentiation protocols have been developed for cartilage and fibrous connective tissues (e.g., tendon), the signals that regulate the induction and differentiation of human iPSCs towards the annulus fibrosus fate remain unknown. Here, we screened a number of candidate factors (and their combinations) and assessed the transcriptomic signatures of key signaling factors involved in embryonic AF development and differentiated function. The transcriptional signatures of treated cells were compared to those of mature human AF cells, and conditions that promoted expression of annulus fibrosus extracellular matrix genes and key transcription factors involved in embryonic AF development were identified. These findings represent an initial approach to guide human induced pluripotent stem cells towards an annulus fibrosus-like fate for cellular delivery strategies.

## Introduction

The intervertebral disc (IVD) is a soft tissue that resides between the bony segments of the spine and is essential for bearing the high loads that arise with activities of daily living. This tissue achieves its function through its highly specialized microarchitecture and biochemical constitution^1^. The disc is a composite tissue that enables motion in six degrees of freedom through its compression-resistant nucleus pulposus (NP) core and its surrounding elastic, high tensile strength annulus fibrosus (AF). Given its demanding loadbearing role, the IVD is often injured, with signs of injury often seen in childhood and in young adults^2^. These insults accumulate over time due to the tissue’s inability to self-repair, leading to severe pathologies such as disc degeneration and herniations later in life.

IVD herniations are one of the most common causes of low back pain – the leading cause of disability worldwide^3^. Disc herniations arise when AF lesions become large enough for migration of NP tissue beyond the anatomical boundaries of the disc. This can result in the compression of adjacent nerves, resulting in debilitating pain and numbness in symptomatic cases^4^. Current clinical strategies for the treatment of symptomatic disc herniations involves resection of the herniated tissue to relieve nerve impingement. However, in such procedures, the AF tear is not repaired, leaving behind a compromised tissue structure that fails to heal and that is predisposed to progressive and repeated herniation at that site. Indeed, the incidence of re-herniation can reach 25% in some instances^5^.

Strategies to repair the AF, including the delivery of growth factors, drugs, and biomaterials, have been explored but have shown limited benefits due to rapid drug clearance, limited molecule diffusion, and a lack of endogenous cell engagement and recruitment^6–8^. The AF is a dense fibrocartilaginous tissue that is formed early during embryonic development by highly anabolic cells that secrete and organize specialized matrix into a functional structure^9,10^. The inner region of the AF accumulates glycosaminoglycans and type II collagen while the outer region is rich in type I collagen and elastin. Once the tissue matures, resident AF cells decrease protein production and only a few cells are left inhabiting the fully formed dense matrix^11,12^. These cells have limited biosynthetic abilities, and are further diminished with injury-induced apoptosis^13^. Therefore, cell therapies that deliver specialized exogenous cells could enhance the restoration of the AF post-herniation.

Cellular delivery strategies for the AF remain underdeveloped due to a lack of AF-specific markers and an optimal cell type for tissue repair. While mesenchymal stromal cells and adipose-derived stem cells have been used in previous attempts, the ability of these cells to take on an AF-specific phenotype is not yet proven. Furthermore, some reports of *in vivo* delivery of these cells resulted in the formation of osteophytes ^14^, highlighting the importance of cell priming and specialization before *in vivo* delivery.

Human induced pluripotent stem cells (iPSCs) have gained widespread use for cellular therapies due to their infinite cell-renewal ability and their potential to differentiate into all three embryonic germ layers through sequential developmentally guided specification. To date, the differentiation of human iPSCs towards the AF fate has not been attempted. While differentiation approaches for NP cells have been reported^15–17^, these have limited application for the differentiation of AF cells due to differences in embryonic lineage (notochord vs. paraxial mesoderm, respectively)^10^. AF cells are specified from mesenchymal progenitors that, also give rise to cartilage, ligament, and tendon during embryonic development^18–20^. Although the iPSC differentiation strategies for cartilage and tendon can be used as guiding principles, the molecular cues that drive human AF specification have not been defined.

During embryonic development, the paraxial mesoderm undergoes segmentation into somites that are dorsoventrally patterned into the dorsal somite (dermomyotome) and the ventral somite (sclerotome)^10,21^. The sclerotome gives rise to the axial skeleton, including bone, cartilage, and fibrocartilaginous tissues. Sclerotome-derived mesenchymal progenitors undergo condensation to form the AF around the NP that is concomitantly forming through notochord segmentation and expansion^10^. A number of molecular factors have been implicated in AF formation. This includes transforming growth factor β (TGF-β), which is indispensable for the formation, maintenance, and growth of the IVD^22,23^. Insulin-like growth factors (IGFs) have also been implicated in skeletal growth and development^24^. Although less is known about their precise role during IVD development, they do stimulate proliferation and extracellular matrix (ECM) synthesis in IVD cells^25,26^. When combined with platelet-derived growth factor (PDGF), IGF reduces apoptosis in human AF cells^25^ and, in concert with TGF-β, increases the production of collagen and other ECM proteins^27^. Moreover, the combination of PDGF and TGF-β directs sclerotome cells towards a fibroblast fate resembling the AF morphology^28^. Another factor, connective tissue growth factor (CTGF), is involved in several cellular functions and is important in skeletal development. During early embryogenesis, CTGF is highly expressed in the somites and notochord, and remains expressed in mature tissues that arise from these embryonic structures^29^. Similarly, sonic hedgehog (SHH) has been implicated in regulating notochord patterning and IVD development^30^. Although these signaling factors have been implicated in IVD development, the effect of these factors (individually or in combination) for the differentiation of AF cells remains unknown.

In this study, we explored the changes induced by developmental signals, independently or in concert, in the differentiation of human iPSCs towards AF-like cells. Due to the lack of molecular markers unique to AF cell population, AF ‘fate’ was assessed by the expression and production of proteins characteristic of AF tissue. Developmental signals were screened by treating iPSC-derived sclerotome cells and assessing gene expression changes at different time points of induction. TGF-β3, in combination with PDGF-BB, CTGF, or IGF-1, induced an upregulation of key AF ECM genes. The synergistic effects observed were validated by using three distinct iPSC lines and by assessing the production of upregulated ECM proteins of interest. Finally, to conduct a broader analysis of the transcriptomic shifts elicited by each factor combination, and to compare genetic profiles of treated cells to mature human AF cells, a 96.96 Fluidigm gene expression array, spanning ECM-related genes, differentiation-specific markers, and genes involved in several cellular processes, was applied. Principal component analysis was employed to identify the transcriptional signatures of each cell population and treatment group in comparison to native AF cells. Together, these studies represent the first step towards developing a differentiation method for iPSC-derived AF cells for future use in cell-based repair of the AF following disc herniation.

## Results

### Differentiation of human iPSCs towards sclerotome

To begin this process, we first differentiated human iPSCs towards a sclerotome fate, using a protocol slightly modified from Adkar et al. Successful sclerotome phenotype was verified in three distinct cell lines (AICS-0061-036, CHOP WT10.2, and CHOP WT4.3) (**Supplementary Figure S1**) ^31^ via gene expression analysis for key stage-specific transcription factors and markers via quantitative RT-PCR (**Supplementary Figure S1**). As expected, over the course of differentiation, there was a gradual decrease in the expression of the pluripotency marker POU domain, class 5, transcription factor 1 (POU5F1), also known as Oct-4 (**Fig. 1a**). After 24 hrs. of anterior primitive streak (APS) induction, the expression of one APS marker mix paired-like homeobox (MIXL1) gene surged and rapidly decreased with the start of paraxial mesoderm (PM) induction. The t-box transcription factor T (TBXT) gene, associated with primitive streak and mesoderm differentiation, was upregulated during the APS and PM stages. Platelet derived growth factor receptor α (PDGFRα), an early mesoderm marker (mainly for paraxial mesoderm), showed a sustained gradual increase from PM to SCL stages. Finally, the sclerotome marker gene SRY-box 9 (SOX9) showed peak expression at the SCL stage after 72 hrs. of sclerotome induction (**Fig. 1a, Supplementary Figure S1**).

**Figure 1.**
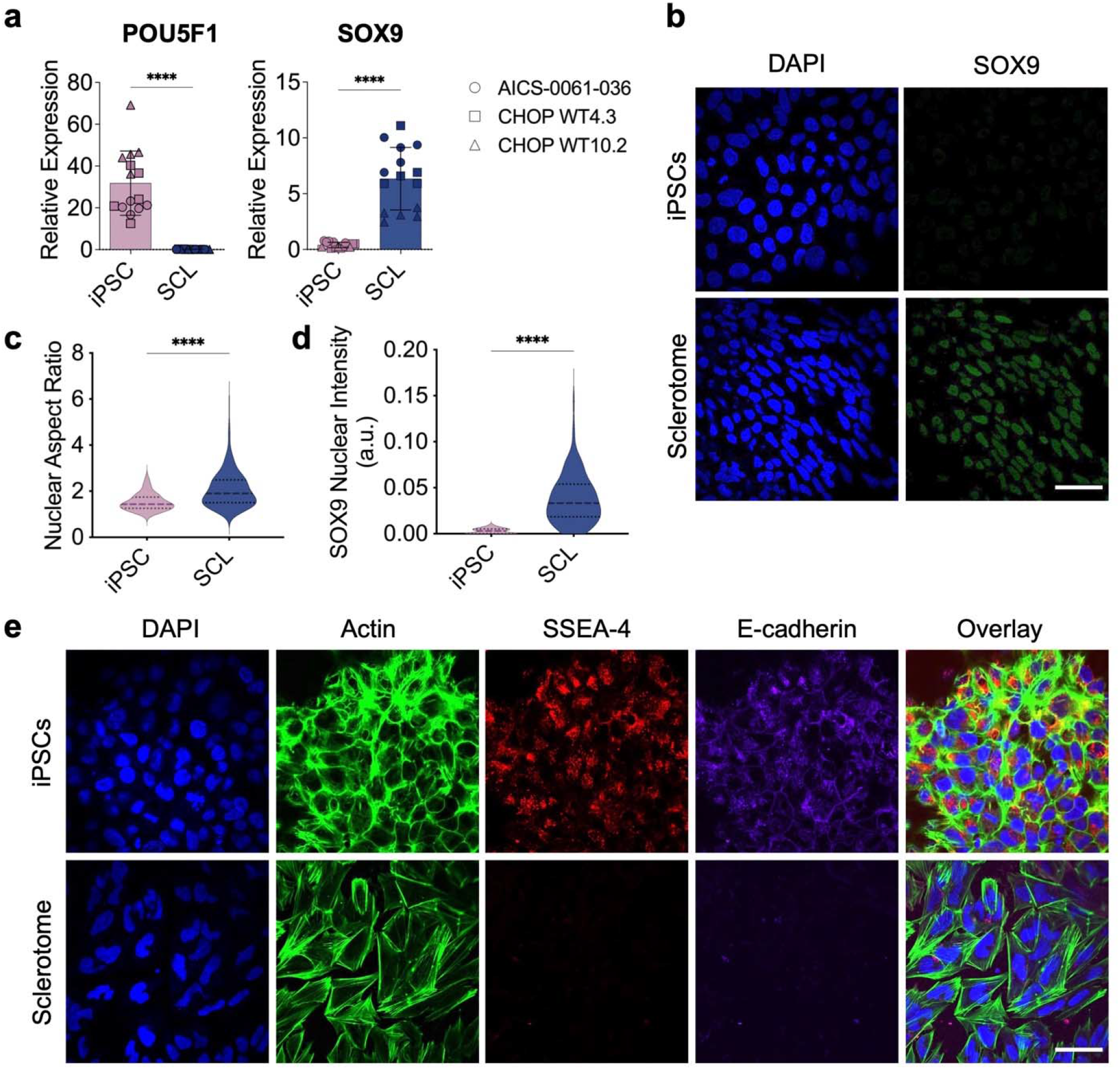
Directed differentiation of human iPSCs towards the sclerotome fate. (a) POU5F1 and SOX9 gene expression at the iPSC and sclerotome (SCL) stages of differentiation for three iPSC cell lines normalized to housekeeping gene expression (n=4/cell line; mean ± s.d.). (b) Representative immunofluorescence imaging for Sox9 intranuclear staining at the iPSC and SCL stages of differentiation (scale: 50 μm). (c) Cell nucleus aspect ratio and (d) Sox9 nuclear intensity measured from immunofluorescence images for iPSC and SCL cells (n>1,000 cells). (e) Representative immunofluorescence imaging for cell nuclear morphology (DAPI), actin filaments (phalloidin), Ssea-4 pluripotency marker, and E-cadherin for iPSC and SCL stages of differentiation (scale: 50 μm). Statistical significance denoted by * p-value < 0.05, ** < 0.01, *** < 0.001, **** < 0.0001.

To further validate the generation of sclerotome cells from human iPSCs, the presence of intranuclear Sox9 was verified via immunofluorescence staining. After 72 hrs. of sclerotome induction, there was an increased intensity of nuclear Sox9 compared to uninduced iPSCs (**Fig. 1b, Fig.1d**). Sclerotome cells also developed an elongated nuclear morphology with greater nuclear aspect ratios compared to iPSCs, characteristic of cellular differentiation (**Fig. 1c**). These changes in nuclear morphology were accompanied by a shift from cortical F-actin to the development of robust F-actin stress fibers (**Fig. 1e**). Furthermore, the sclerotome cells lost expression of the pluripotency marker SSEA-4 and showed a reduction in cell-cell contacts as evidenced by a decrease in E-cadherin staining, corroborating the shift from a pluripotent to a differentiated state (**Fig. 1e**).

### Candidate screening reveals synergistic role of factors in combination with TGF-β to drive annulus fibrosus-like fate

After validating the iPSC-derived sclerotome cells, factors present during the embryonic development of the IVD and AF were investigated with respect to their ability to promote induction of an AF-like fate. TGF-β3, CTGF, PDGF-BB, IGF-1, or the Hedgehog pathway activator, Purmorphamine (Pu) were added independently or in combination with TGF-β3 (due to its instrumental role in IVD morphogenesis) (**Fig. 2a**). The expression of genes for major AF extracellular matrix (ECM) proteins including type I collagen (COL1A1), type II collagen (COL2A1), elastin (ELN), and aggrecan (ACAN) was compared among groups after 7 or 14 days of treatment with different combinations. All genes assessed showed peak expression after 14 days of treatment compared to 7 days (**Fig. 2b-c**). After 7 days of induction, the group treated with TGF-β3 + PDGF-BB (+TP) showed significantly higher expression of COL1A1, ELN, and ACAN compared to the untreated control cells (Ctrl) and other treatment groups. By 14 days of exposure to factors, the dual-factor combination groups that included TGF-β3 (i.e., +TP, +TC, and +TI), all showed significantly greater expression of COL1A1, ELN, and ACAN compared to the Ctrl group, and trended towards greater expression of COL2A1. Interestingly, combining TGF-β3 with Purmorphamine (+TPu) did not show this effect. Furthermore, the addition of the Hedgehog pathway activator in the triple factor combinations (+TPPu, + TCPu, and +TIPu) decreased the expression of these ECM genes by 14 days compared to the double factor combinations that did not include Purmorphamine (+TP, +TC, and +TI) (**Fig. 2c**). Based on these findings, the Hedgehog pathway activation via the addition of Purmorphamine was not pursued further.

**Figure 2.**
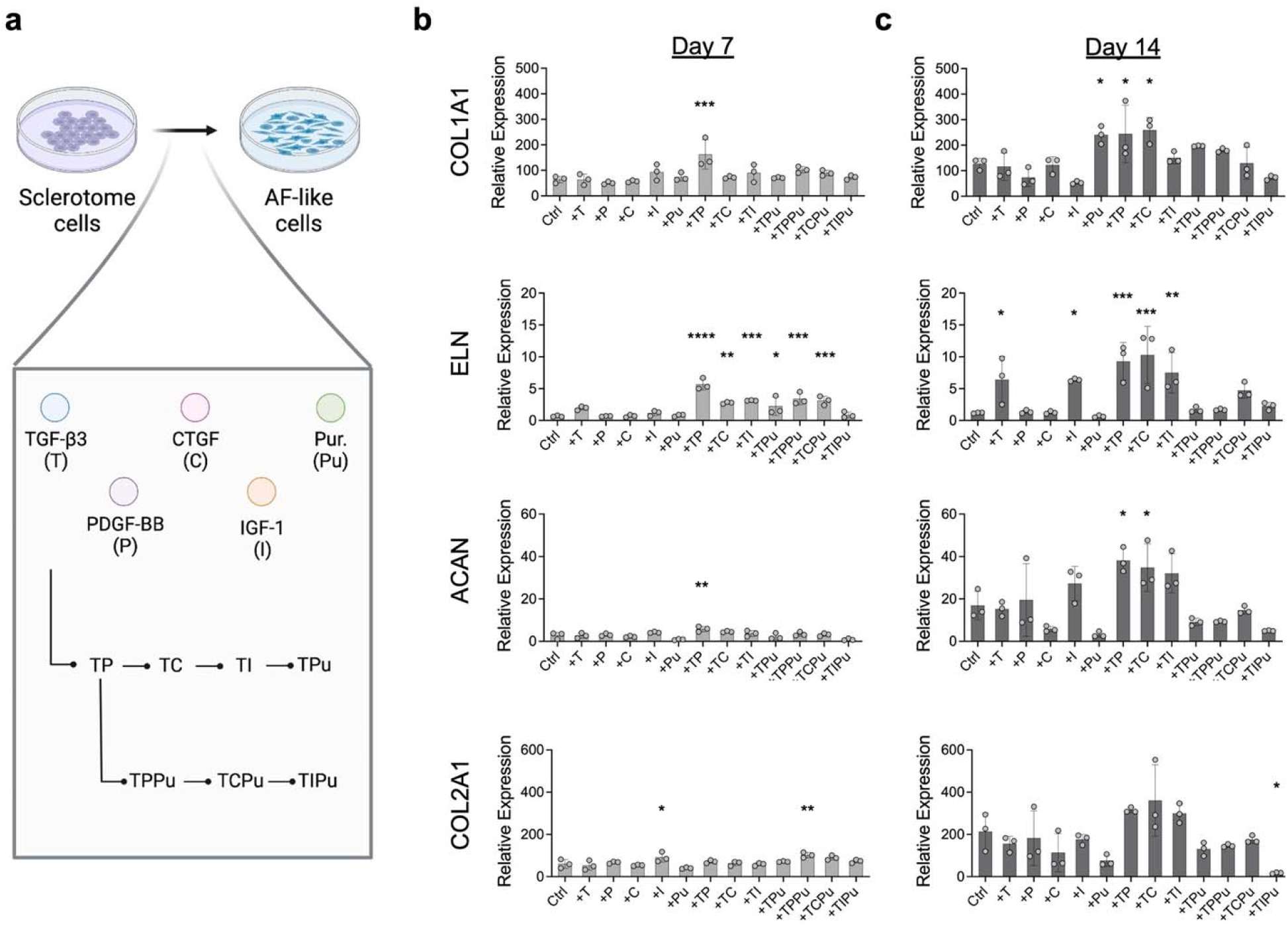
TGF-β in combination with PDGF-BB, CTGF or IGF-1 drives the upregulation of major AF ECM genes. (a) Schematic of the factors tested to drive sclerotome cells towards the annulus fibrosus fate. (b) Gene expression for major annulus fibrosus extracellular matrix protein genes, COL1A1, ELN, ACAN, and COL2A1, after 7 or 14 days of exposure to differentiation factors, normalized to housekeeping gene expression (n=3; mean ± s.d.). Statistical significance compared to Ctrl denoted by * p-value < 0.05, ** < 0.01, *** < 0.001, **** < 0.0001.

To verify that the double factor treatments led to comparable effects across iPSC cell lines, +TP, +TC, and +TI treatments were assessed using 3 distinct lines and compared to Ctrl. An additional group in which these factors were combined (+TPCI) was added to this analysis. After 14 days of treatment, expression of COL1A1, COL2A1, ELN, and ACAN was greater for all treatment groups compared to Ctrl, across all cell lines (**Fig. 3a**). The +TP group resulted in significantly higher expression of COL2A1 and ACAN, while ELN was significantly higher than Ctrl with +TC or +TI treatment after 14 days (**Fig. 3a**). The combination of TGF-β3, PDGF-BB, CTGF, and IGF-1 did not have a synergistic effect on the expression of any these key AF ECM genes. To assess whether changes in gene expression translated to the deposition of matrix, immunofluorescence staining for type I collagen and elastin – the major structural proteins of the outer AF – was performed after 14 days of treatment (**Fig. 3b**). Compared to Ctrl, induced cells showed a greater distribution and density of collagen and elastin deposition, providing evidence of AF-like matrix elaboration (**Fig. 3b**).

**Figure 3.**
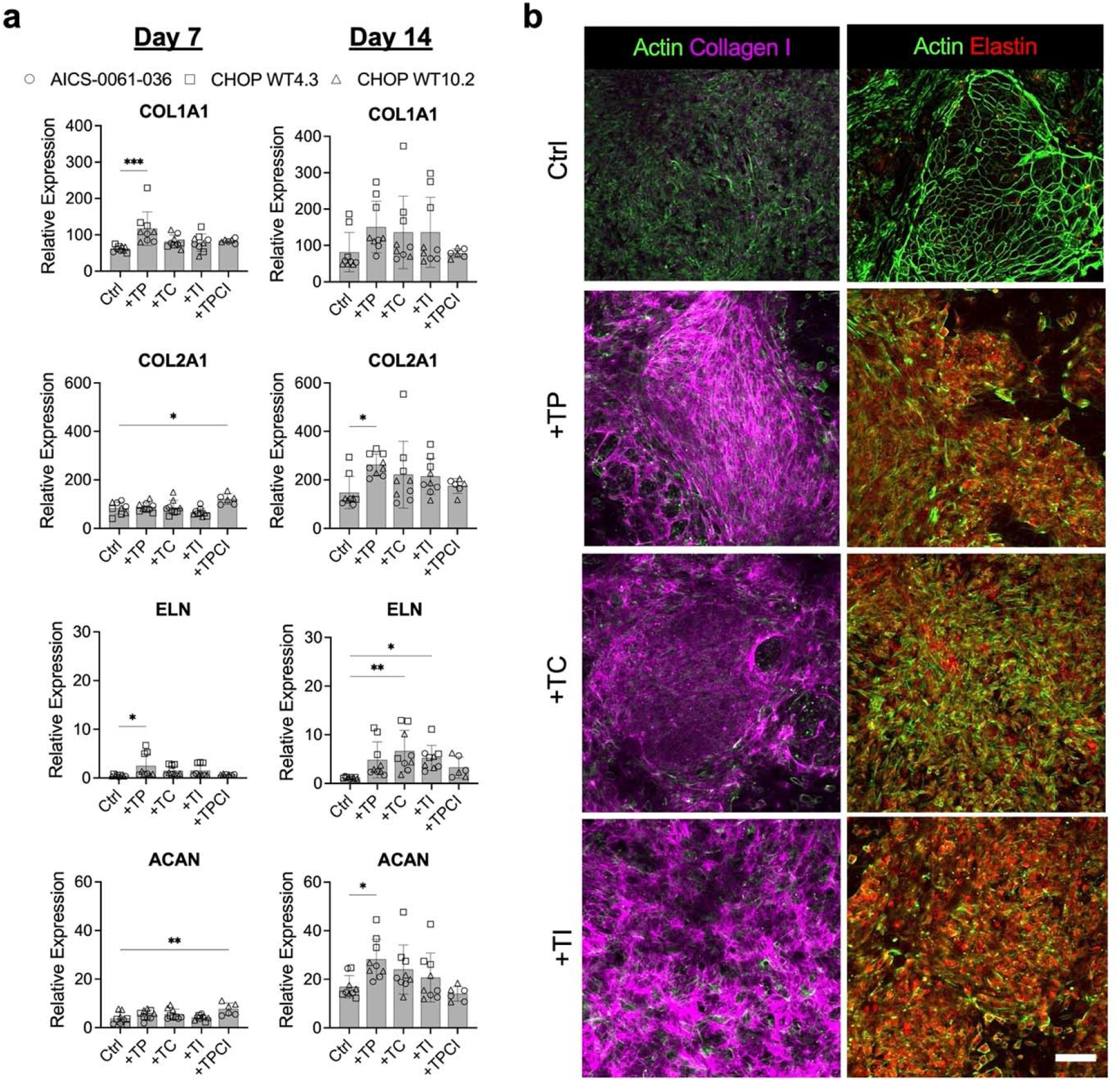
Upregulated expression and production of major annulus fibrosus extracellular matrix proteins with top performing factor combinations. (a) Gene expression for major annulus fibrosus extracellular matrix protein genes COL1A1, ELN, COL2A1, and ACAN after 7 or 14 days of exposure to top performing differentiation factor combinations, normalized to housekeeping gene expression for three different iPSC cell lines (n=3/cell line; mean ± s.d.). (b) Immunofluorescence staining for type I collagen and elastin, co-stained for F-actin (phalloidin) after 14 days of exposure to factor combinations (scale: 100 μm). Statistical significance compared to Ctrl denoted by * p-value < 0.05, ** < 0.01, *** < 0.001, **** <0.0001.

### Fluidigm Gene Expression Array for the Comparison of Factor Treatment Effects

To conduct a broader analysis of the gene expression differences between human AF cells and iPSC derived SCL cells that were induced towards AF differentiation, a Fluidigm 96.96 gene expression array was carried out. This enabled simultaneous comparison of adult AF, SCL, and +TP, +TC, +TI, +TPCI groups after 14 days of treatment (**Supplementary Figure S2**). 93 genes, spanning collagens, proteoglycans and glycoproteins, ECM-remodeling factors, differentiation specific markers, cell-cell and cell-ECM interaction proteins, and mechanotransduction mediators, were included in the qPCR array (**Supplementary Table S4**). TBP, 18S, and RPS17 were used as housekeeping genes.

From these data, we first carried out principal component analysis (PCA) of ΔC_t_ values for healthy human adult AF vs. SCL cells to identify the genetic signatures of each cell population (**Fig. 4a, Supplementary Figure S3**). PC1 captured 74.5% of the total variance while PC2 captured 9.8%, together representing 84.3% of the variance between the groups (**Fig.4b**). PC1 scores were significantly different between groups, with AF scores being higher than SCL scores, while no differences in PC2 scores were observed (**Fig. 4c**). Several genes involved in the establishment and maintenance of AF ECM (TIMP1, FN1, COL6A1, SPARC, DCN) were among the top 8 genes with highest PC1 loading values. Other top scoring PC1 loading genes included THY1, ACAN, LUM, CA12, TGFBR2, COMP, ITGA5, COL12A1, GDF5, PAX1, PXN, LGALS3, MYH9, COL1A1, ROCK2, IGFBP7, PIEZO1, COL1A2, CTGF, TGFB1, and ITGB5 (**Supplementary Table S5**). Meanwhile, several of the lowest PC1 loading value genes (TBXT, TBX6, MSGN1) represented established mesoderm markers, characteristic of SCL cells (**Fig. 4d**). This analysis thus provided a clear set of markers to define genetic profiles for each cell population.

**Figure 4.**
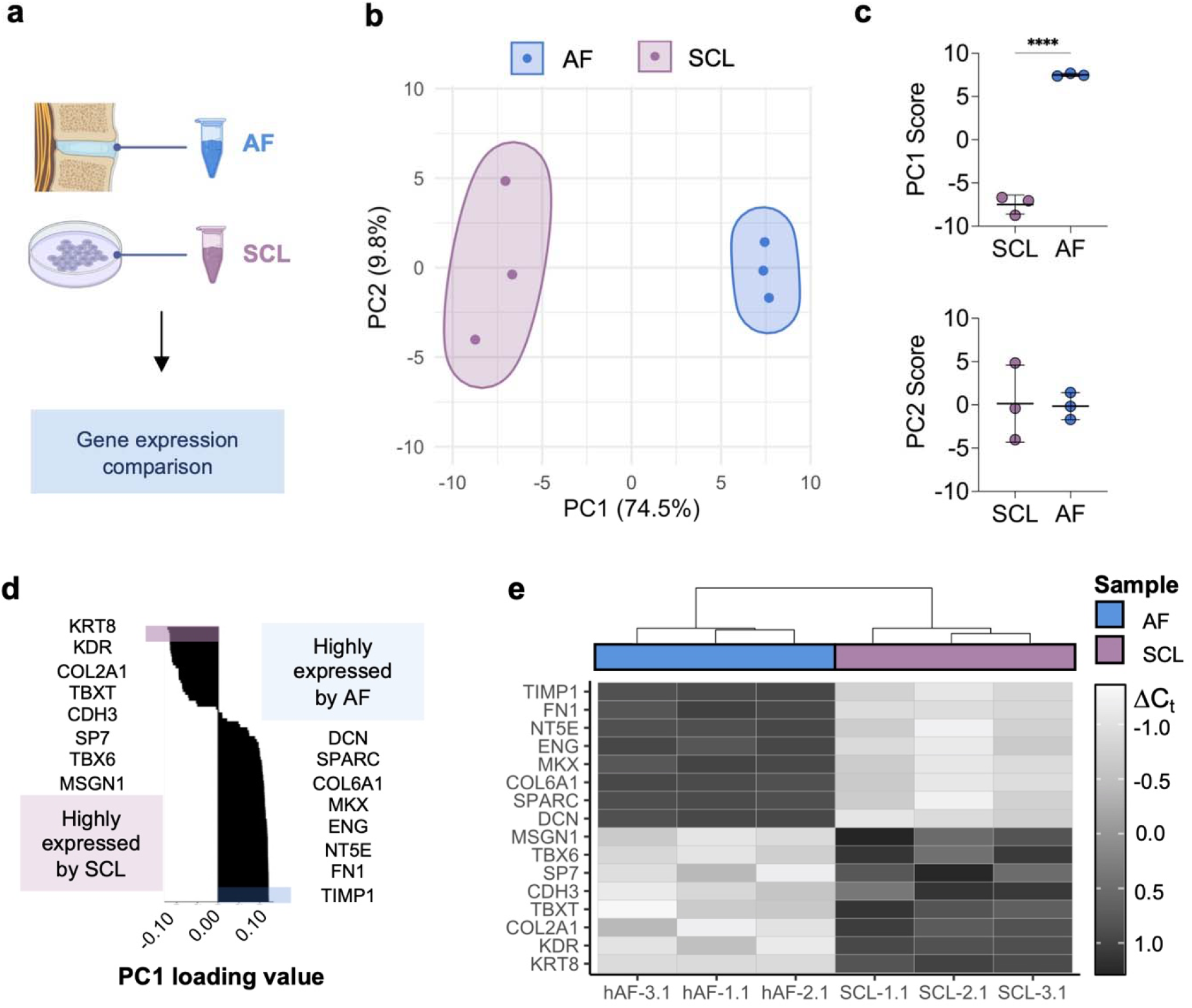
Principal component analysis of SCL and AF cells reveal differences in Fluidigm gene expression array profiles. (a) The gene expression patterns of AF cells from 3 healthy human donors and SCL cells differentiated from 3 human iPSC cell lines were compared. (b) Principal component analysis (PCA) of the data represented by mapping PC1 and PC2 scores for SCL and AF groups. Percent of described total variance by each PC is listed in parentheses. Concentration ellipses for each group are demarcated via clustering. (c) (Top) PC1 scores and (Bottom) PC2 scores for SCL and AF groups (n=3/group; mean ± s.d.). (d) Ranked PC1 loading values for genes included in the Fluidigm gene expression array, listing the 8 highest and 8 lowest valued genes. For a full list of ranked PC1 loading values, see **Supplementary Table S5**. (e) Heatmap of ΔC_t_ values for genes listed in (d) with hierarchical clustering based on the 93 genes included in the gene expression array with color coding by sample type. For the heatmap showing all 93 genes used in the cluster analysis, see **Supplementary Figure S3**. Statistical significance denoted by * p-value < 0.05, ** < 0.01, *** < 0.001, **** <0.0001.

Next, to assesses the genetic differences between different treatment groups, PCA was conducted using ΔC_t_ values for AF, SCL, and SCL cells treated for 14 days with factor combinations (TP14, TC14, TI14, and TPCI14) across the 93 genes analyzed with the Fluidigm gene expression array (**Supplementary Figure S4**). A heatmap of the processed data revealed first-level clustering of the day 14 treatment groups with SCL cells, separated from AF cells. This indicates that, after 14 days of induction, treated cells continued to show greater gene expression similarities to SCL cells compared to adult human AF cells. Second-level clustering revealed differences between SCL cells and all day 14 treatment groups.

Interestingly, the iPSC cell line AICS-0061-036 (AICS) showed cluster separation from the CHOP WT4.2 (WT4.2) and CHOP WT10.3 (WT10.3) cell lines. Further assessment of the global gene expression patterns showed marked differences across several genes between AICS and the other two cell lines. Specifically, all AICS cells, regardless of treatment type, showed lower expression of key genes that were highly upregulated by adult AF cells compared to WT4.2 and WT10.3-treated cells. However, this marked difference in gene expression was not apparent at the SCL stage, indicating the cell line-dependent differences arose in later stages of differentiation with extended factor treatment (**Supplementary Figure S4**).

From the overall analysis, PC1 and PC2 scores captured 62.1% of the variance between the groups assessed. Plotting of PC1 and PC2 scores for each sample showed overlap between all day 14 treatment groups (TP14, TC14, TI14, and TPCI14), with all populations lying between SCL and adult AF cells (**Supplementary Figure S5**). Further inspection of the PC1 and PC2 score differences among treatment groups revealed no significant differences after 14 days of induction with the different factor combinations (**Supplementary Figure S5**), suggesting that the differences between treatment groups did not induce drastic differences in gene expression. Assessment of the 8 highest and 8 lowest scoring genes identified from the PCA of adult AF and SCL cells showed increased expression of genes characteristic of AF cells with 14 days of induction and a reduction in expression of genes highly expressed at the SCL stage across all treatments (**Supplementary Figure S5**). However, the AICS cell line showed lower expression of AF-specific genes compared to WT4.2 and WT10.3, regardless of factor combination.

To further explore the effects of treatment, we compared genetic profiles of the TP14 group to those of SCL and adult AF cells. TP14 was chosen due to the significantly increased expression of COL2A1 and ACAN compared to other treatment groups, in addition to the physical manifestation of COL1A1 and ELN in IF staining (**Fig. 3a**). PCA of AF, SCL, and TP14 is shown in **Fig. 5a**. PC1 score comparisons, which captured 52.8% of the variance, revealed significant differences between all groups (**Fig. 5b**). PC1 scores gradually increased as a function of differentiation progression, increasing from SCL to the adult AF stage. Meanwhile, the PC2 score, which captured 17.7% of the variance, only revealed differences between the SCL and TP14 group. TP14 and AF cells did not have significantly different PC2 scores (**Fig. 5b**).

**Figure 5.**
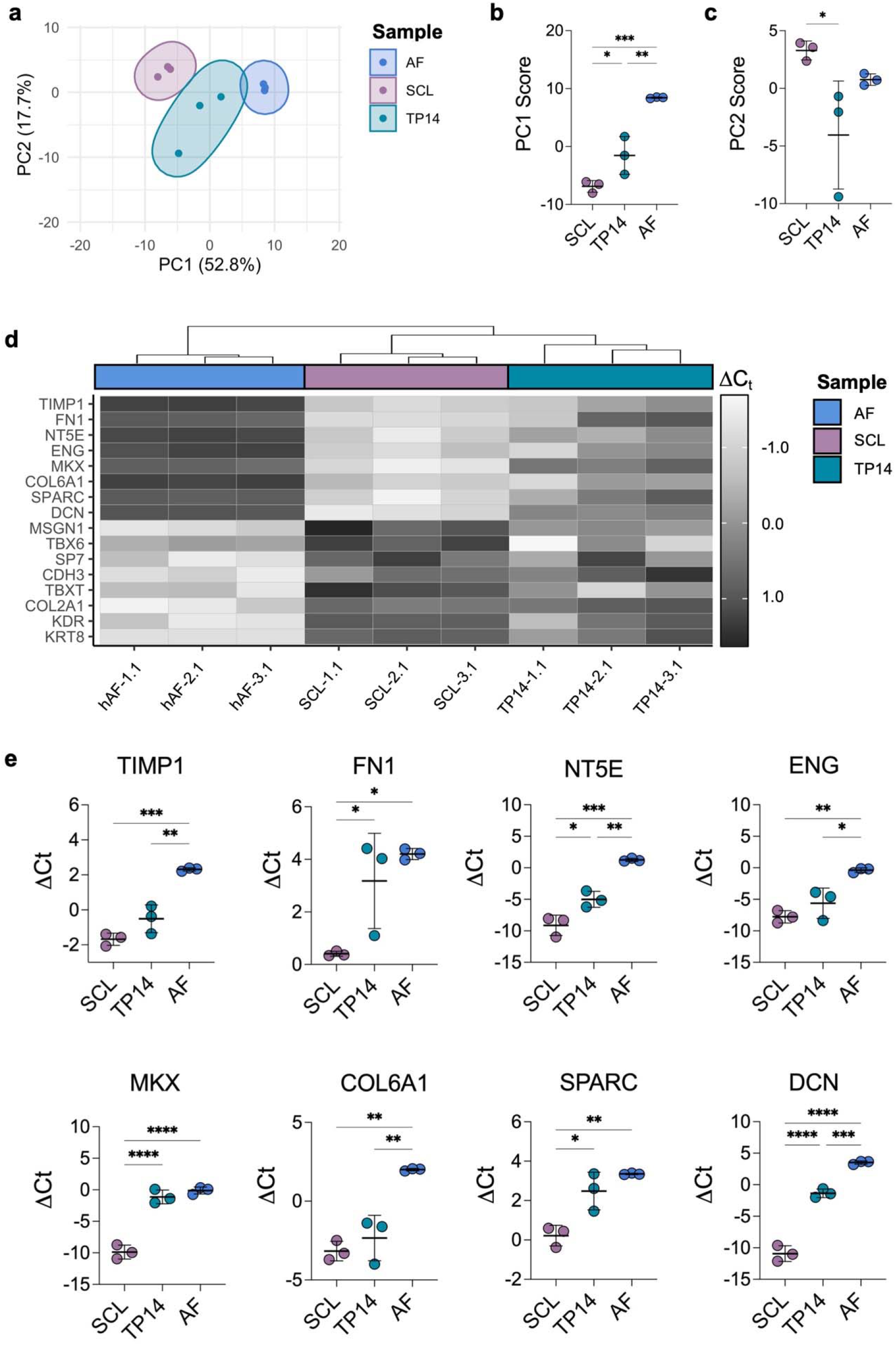
Treatment of SCL cells with factors involved in AF embryonic development drive changes in gene expression towards the expression patterns of adult AF cells. (a) PCA of the data represented by mapping PC1 and PC2 scores for SCL, AF, and TP14 groups. Percent of described total variance by each PC is listed in parentheses. Concentration ellipses for each group are demarcated via clustering. (b) PC1 scores and (c) PC2 scores for SCL, AF, and TP14 groups (n=3/group; mean ± s.d.). (d) Heatmap of ΔC_t_ values for the 8 highest and 8 lowest valued genes identified from the AF vs. SCL PCA, with hierarchical clustering based on the 93 genes included in the gene expression array with color coding by sample type. For the heatmap showing all 93 genes used in the cluster analysis, see **Supplementary Figure S4**. (e) ΔC_t_ values for genes with highest PC1 score in AF vs. SCL PCA (highest expression shown by AF cells) (n=3/group; mean ± s.d.). Statistical significance denoted by * p-value < 0.05, ** < 0.01, *** < 0.001, **** <0.0001.

Assessment of the 8 genes uniquely highly expressed by AF cells or SCL cells, respectively, revealed a shift towards the genetic profile of AF cells with the addition of TGF-β3 and PDGF-BB for 14 days (TP14) (**Fig. 5d, e**). Specifically, addition of these factors increased the expression of FN1, MKX, a key transcription factor involved in AF development, and SPARC to levels expressed by mature adult AF cells (**Fig. 5e**). Other genes highly expressed by mature AF cells such as TIMP1, NT5E, ENG, COL6A1, and DCN also showed increases in expression with treatment. Concomitantly, genes highly expressed by SCL cells were downregulated with treatment, with TBX6 and TBXT reaching expression levels apparent in mature AF cells (**Supplementary Figure S6**). Other genes highly expressed by SCL, such as MSGN1, SP7, KDR, and KRT8, were reduced to a lesser extent by treatment.

Expression of ECM-related genes such as COL1A1, COL12A1, ACAN, PRG4, LUM, COMP, and FMOD was dramatically upregulated with treatment, in some cases reaching levels comparable to mature AF cells (**Fig. 6a**). Other genes involved in the TGFB signaling pathway such as TGFB3, and IGF-related genes (IGF-1, IGFBP5, and IGFBP7) showed expression changes towards levels measured in adult AF cells with treatment (**Fig. 6b**). The increase in expression for AF ECM proteins in treated cells was accompanied by a reduction in the expression of genes involved in early embryonic development, including PODXL, FOXC2, and proteins involved in cell-to-cell adhesion, namely CDH4 and CDH5 (**Fig. 6c**). Other genes involved in diverse cellular processes, such as PIEZO2 and GDF5, showed expression shifts towards AF levels with treatment (**Fig. 6d**). Interestingly, expression of the Scleraxis transcription factor (Scx) gene, SCX, required for AF and tendon development, was markedly upregulated with treatment, surpassing expression levels measured in the adult AF (**Supplementary Figure S7**). Similarly, the expression of TNMD and ADAMTS6 surpassed that of mature AF cells with treatment, while PAX1 expression decreased with treatment (**Supplementary Figure S7**). These results demonstrated that treatment of SCL cells with a combination of TGF-β3 and PDGF-BB for 14 days differentiated cells away from SCL towards an anabolic state characteristic of fibrocartilaginous cells.

**Figure 6.**
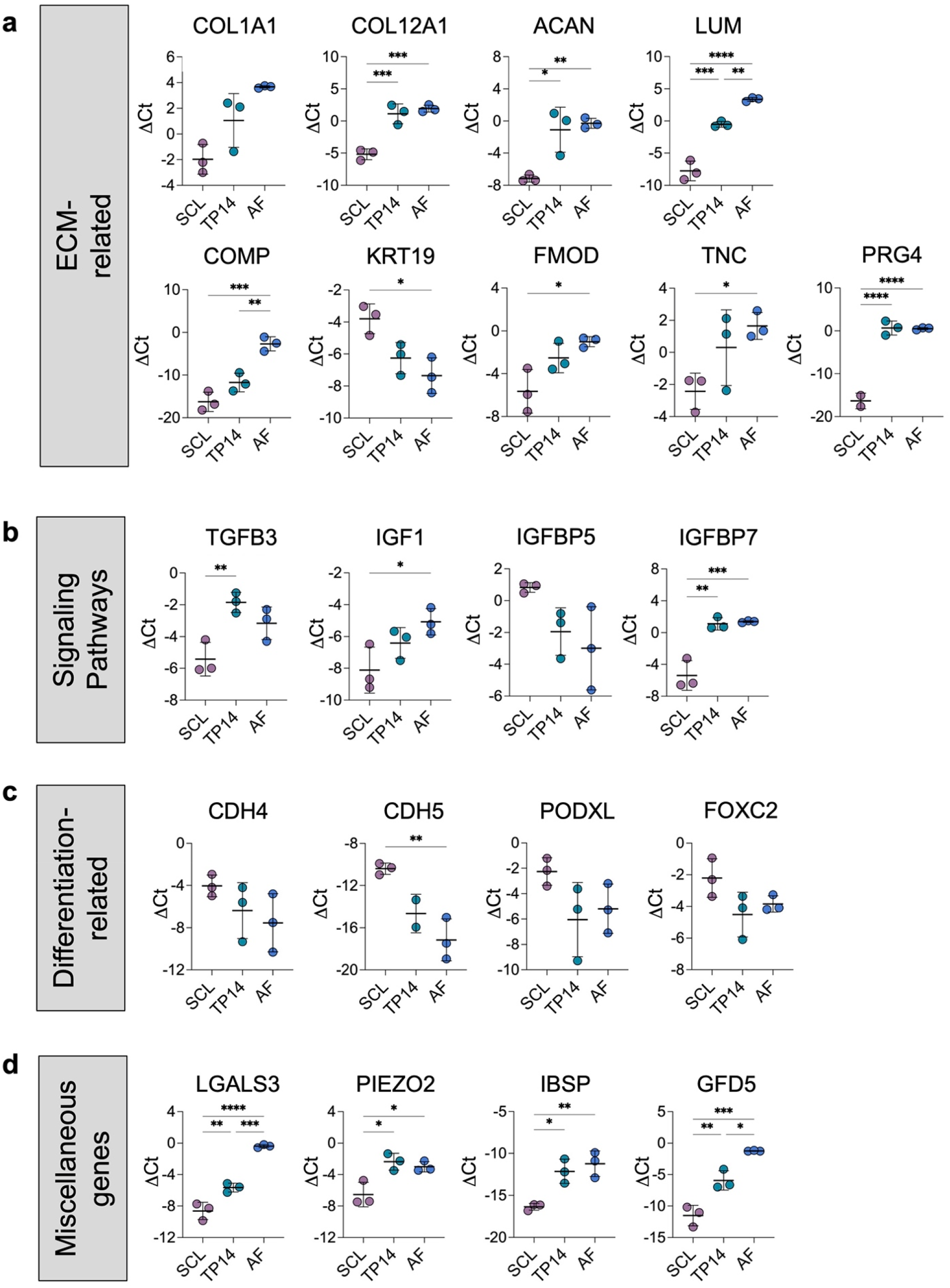
Factor treatment drives changes in gene expression towards an anabolic AF-like state. (a) ΔC_t_ values for ECM-related, (b) signaling pathways, (c) differentiation-related, and (d) other genes involved in a variety of cellular processes for SCL, TP14, and AF cells (n=3/group; mean ± s.d.). Statistical significance denoted by * p-value < 0.05, ** < 0.01, *** < 0.001, **** <0.0001.

## Discussion

In this study, we screened developmental molecular signals involved in axial skeletal and IVD development to establish their effects on the induction of an AF-like phenotype in human iPSCs. The impact of TGF-β, PDGF, CTGF, IGF-1, and Hedgehog signaling were investigated individually or in combination with TGF-β, which is known to be indispensable for disc development, maintenance, and maturation^22,23^. Using this screen, the synergy of TGF-β, when combined with other factors, was evidenced at the transcriptomic level. This was marked by a dramatic increase in expression of AF-specific ECM genes that resulted in increased translation of AF-specific ECM proteins. Using a 96.96 Fluidigm gene expression array, the transcriptional signatures of cells before and after treatment were compared to that of mature AF cells. This established that combinations of developmental signals resulted in iPSC-derived sclerotome (SCL) cells to partially shift away from their primitive SCL fate and towards the pro-anabolic mature AF phenotype.

To date, AF healing has only been shown during fetal and neonatal stages, indicating that early AF cells have a unique reparative potential ideal for tissue replenishment and healing^32,33^. Recreating this early AF phenotype from human iPSCs has never been attempted but may represent a promising approach as a cell therapy for disc herniation interventions. Although TGF-β has been identified as a key regulator of IVD formation, the effect of other signals present during AF embryonic development remains unknown. In this study, we demonstrate that TGF-β alone is insufficient to drive the upregulation of AF-specific ECM genes. Similarly, other signals such as PDGF-BB, CTGF, IGF-1, or SHH activation had limited effects when applied independently. However, TGF-β combinations with PDGF-BB, CTGF, or IGF-1 dramatically increased the expression of major AF ECM proteins, which resulted in increased matrix deposition. Surprisingly, broader examination of transcriptomic changes resulting from these signal combinations through a 96.96 Fluidigm gene expression array did not show significant differences between treatment combinations. This indicates that although TGF-β synergizes with PDGF-BB, CTGF, or IGF-1, the synergy remains relatively comparable for the genes assessed. Conducting bulk or single cell RNA sequencing could provide a greater appreciation of the transcriptomic signatures of each cell population.

Even though our analysis did not yield differences between the TGF-β combinations, all treatments drove differences in gene expression towards the AF fate and away from the SCL fate. Analysis of the TP14 group showed a significant pro-anabolic shift in cells, where 14 days of treatment induced the upregulation of major and minor AF-specific ECM proteins involved in embryonic collagen fibrillogenesis^34^. Strikingly, Scx and Mkx, two transcription factors highly involved and necessary for AF development, were significantly upregulated by treatment. SCX expression in treated cells reached levels greater than that of the mature AF cell expression level, in line with the high SCX expression observed in the embryonic AF that is attenuated with maturation^35,36^. Scx-lineage cells have also been implicated in the only reported instances of AF healing and in the proper attachment of ligamentous tissues^33,35,37^. AF healing in neonates is directed by Scx-lineage cells that proliferate and deposit type I collagen to repair AF lesions^33^. Scx-lineage cells that express TNMD and SOX9, two genes upregulated by treatment, have also been implicated in proper AF formation^33,35^. Similar to Scx, Mkx was upregulated with treatment to levels observed in mature AF cells^38,39^. Together, these results highlighted the strong inductive potential of the signal combinations used to direct the AF specification of SCL progenitors.

Although several AF-specific genes were upregulated upon treatment, PCA analysis revealed that treated cells remained closer to SCL cells than to mature AF cells. This may be due the comparison of early embryonic cells to fully formed AF cells obtained from adult donors. Obtaining human embryonic AF tissue would provide the most accurate comparison, but this remains a challenging endeavor. Another potential reason for this difference is the lack of additional induction signals that may complete the differentiation process. Our study explored limited signaling players and did not investigate the induction effects of different concentrations, timing of initial exposure, and exposure duration. In addition to investigating these different aspects of treatment and other morphogen signals, the biophysical cues present during AF development could enhance AF specification. During embryonic development, the AF and surrounding tissues undergo continual growth and stiffening in unison with morphogenetic events^40^. Growth-mediated stresses and strains can dictate tissue segmentation patterns, activate molecular signaling events, and provide structural integrity to tissue structures during key morphogenetic events, playing an indispensable role throughout embryonic development^41,42^. Exploring the interplay between morphogens and biophysical cues could further improve the differentiation strategies employed for the generation of AF cells.

The AF is a tissue with numerous endogenous impediments for repair^6^. Here, we set the stage for the development of iPSC-based cellular therapies for AF repair. Future investigations exploring more complex signals and their interplay with morphogens that guide AF fate induction will enable the development of a reparative AF cell population for the restoration of the AF structure and function. Combining the delivery of these cells with the surgical removal of herniated tissue represents a promising approach to improve repair outcomes and lower recurrent herniation rates after surgical intervention.

## Methods

### iPSC Maintenance and Expansion

The human iPSC line AICS-0061-036 was purchased from the Coriell Institute for Medical Research. The CHOP WT10.3 and CHOP WT4.2 human iPSC cell lines were donated by the Human Pluripotent Stem Cell Core at the Children’s Hospital of Philadelphia. For expansion and maintenance, human iPSCs were cultured in tissue culture plates coated with hESC qualified Matrigel (BD Biosciences, 352277) with feeder-free mTeSR™1 maintenance medium (Stemcell Technologies, 85850). The enzyme-free Gentle Cell Dissociation Reagent (Stemcell Technologies, 100-0485) was used to dissociate iPSCs into cell aggregates for routine passaging.

### iPSC Differentiation

#### Differentiation of human iPSCs towards sclerotome

Human iPSCs were passaged as cell clusters and seeded on tissue culture plates coated with Growth Factor Reduced Matrigel Matrix (Corning, 354230). At ∼70% confluence, cells were fed with mTESR™1 approximately three hours before the start of the differentiation protocol. Washing Medium, composed of 50% IMDM, GlutaMAX™ (ThermoFisher, 31980030) and 50% Ham’s F12 Nutrient Mix (Corning, MT10-080-CV), was used to wash the cells two times before the start of differentiation. Cells were washed with Washing Medium before each differentiation step.

Human iPSCs were induced to differentiate towards the sclerotome fate following a previously established protocol, with slight modifications^31^. The base medium used did not include penicillin/streptomycin and polyvinyl alcohol as reported by Adkar and colleagues. Briefly, iPSCs were guided through a stepwise differentiation approach wherein cells were directed towards the following fates in the respective order: anterior primitive streak (APS) for 24 hrs., paraxial mesoderm (PM) for 24 hrs., early somite (ES) for 24 hrs., and sclerotome (SCL) for 72 hrs. (**Supplementary Fig. S1**). Mesoderm Differentiation Medium, composed of Washing Medium supplemented with 1% Chemically Defined Lipid Concentrate (ThermoFisher, 11905031), 1% Corning ITS™ Premix (Fisher, CB-40350), and 450 μM of 1-thiolglycerol (Sigma, M6145), was used for all differentiation steps, with media changes every day. For each differentiation step, Mesoderm Differentiation Medium supplemented with stage-specific factors, was added to cell cultures as depicted in **Supplementary Fig. S1** (**Supplementary Table S1**). At the end of each differentiation stage, cell samples (n=4/cell line) were collected and frozen for gene expression analysis.

#### Factor screening for induction of annulus fibrosus fate

After the 72 hours of sclerotome induction, cells were washed twice with Washing Medium. Control (Ctrl) wells were fed Mesoderm Differentiation Medium. To screen factors for their ability to promote the AF fate, transforming growth factor beta 3 (TGF-β3) (+T, 10 ng/mL), connective tissue growth factor (CTGF) (+C, 100 ng/mL), platelet derived growth factor BB (PDGF-BB) (+P, 2 ng/mL), insulin-like growth factor 1 (IGF-1) (+I, 5 ng/mL), or the Hedgehog pathway activator, Purmorphamine (+Pu, 2 μM) were added independently or in combination (**Fig. 2a**) (**Supplementary Table S1**) for 7 or 14 days, after which cell samples were either fixed for immunofluorescence staining or collected, pelleted, and frozen at -80°C for gene expression analysis (n=3/cell line).

### Gene Expression Analysis via Quantitative RT-PCR

RNA isolation of digested samples was performed using the Directzol RNA Miniprep kit with DNAse-I treatment to remove trace DNA before RNA elution (Zymo Research, R2050). RNA was quantified via Nanodrop spectrophotometry. cDNA was synthesized using the SuperScript™ IV VILO Master Mix (Invitrogen, 11756050) according to the manufacturer’s protocol. Human genomic DNA, extracted from human cells using the GeneJET Genomic DNA Purification kit (ThermoFisher, K0721), was used to generate a standard curve (300,000-0 copies). Absolute quantitative RT-PCR was run using Fast SYBR™ Green Master Mix (ThermoFisher, 4385618) for 40 cycles with validated custom designed primers (**Supplementary Table S2**), using a genomic DNA standard curve. Changes in gene expression, reported as relative expression, were quantified by normalizing copy numbers for each gene of interest in each sample to the copy numbers of the TBP housekeeping gene in the respective sample (**Supplementary Table S2**).

### Immunofluorescence Staining and Imaging

#### Staining

For immunofluorescence staining, iPSCs were seeded on μ-Slide 8 well polymer coverslip chamber slides (Ibidi, #80826). Differentiation steps were followed as described, including coating wells with Growth Factor Reduced Matrigel before cell seeding. At the defined time points, cells were washed 3x with PBS, after which they were fixed with 4% paraformaldehyde for 20 mins. at room temperature (RT). After fixation, the cells were washed 3x with PBS. For intracellular staining, cells were permeabilized with 0.1% Triton X-100 solution in PBS for 10 mins. at 4°C, after which they were rinsed 3x with PBS. For both intracellular and extracellular staining, blocking was performed by adding blocking solution composed of 3% bovine serum albumin in PBS for 1 hr. at RT. After blocking, cells were washed with PBS 3x, and primary antibodies were added overnight at 4°C (**Supplementary Table S3**). Cells were then washed with PBS 3x and secondary antibodies, phalloidin, and/or DAPI were added for 1 hr. at RT, after which cells were washed 3x with PBS and imaged.

#### Confocal Imaging

Confocal imaging was performed using a Nikon A1R+ confocal microscope. For SOX9 intranuclear staining, cells counterstained with DAPI, and for cells co-stained for F-actin, SSEA-4, and E-cadherin, images were obtained at the mid-plane using 60X magnification. For visualization of type I collagen and elastin, and for imaging of cells co-stained for F-actin, images were obtained at 20X magnification.

#### Nuclear Morphology and Sox9 Intranuclear Intensity Measurements

Nuclear aspect ratio and levels of SOX9 intranuclear fluorescence intensity were assessed using CellProfiler, ver 3.1.9. Nuclei were viewed using DAPI and were segmented based on a minimum cross entropy algorithm with nuclei excluded below 20 and above 60 pixels. Nuclei that were touching the image boundary were also excluded. Major and minor axes were measured, and their ratio (long to short axis) was calculated and is presented as the nuclear aspect ratio (n>1000 cells/sample). SOX9 average fluorescent intensity was measured on a per-nucleus basis using nuclei identified by minimum cross entropy algorithm (n>1000 cells/sample).

### Fluidigm 96.96 Gene Expression Assay

#### Sample Sourcing and Preparation

One sample for each of the three iPSC lines at SCL stage and day 14 of culture for the +TP, +TC, +TI, and +TPCI groups (TP14, TC14, TI14, and TPCI14, respectively) were collected, pelleted, and frozen. Primary human AF cells obtained from healthy human tissue were purchased from Articular Engineering. AF cell lines AF26 (30-year-old donor), AF1356 (26-year-old donor), and AF1363 (59-year-old donor) were used at passage 1.

#### RNA Isolation, cDNA Preparation, and Preamplification

RNA was isolated from the 45 samples (see above) as previously described with DNase-I treatment before RNA elution (see above). cDNA was synthesized using the SuperScript™ IV VILO Master Mix with ezDNase™ Enzyme (Invitrogen, 11766050), with a second DNase digestion to remove gDNA according to manufacturer’s protocol. No reverse transcriptase controls were generated from pooled iPSC, SCL or AF cells by not including reverse transcriptase for cDNA synthesis. Generated cDNA for the 48 samples was pre-amplified for 14 cycles using the specific 96 gene targets. For this, a primer pool containing the 20X TaqMan primers for the gene targets was used (**Supplementary Table S4**), consisting of the primers diluted 1:100 (0.2X) in DNA suspension buffer (10 mM Tris, pH8.0, 0.1 mM EDTA). 2 μL of each cDNA sample was combined with 1.25 μL of pooled diluted TaqMan probes, 1 μL of Fluidigm Preamp Master Mix (Fluidigm, 100-5580), and 0.75 μL of water for a combined 5 μL reaction volume. The resultant pre-amplified cDNA was diluted 1:5 in the DNA suspension buffer.

#### Fluidigm Gene Expression Chip & Data Post-Processing

qPCR of samples (n=3 cell lines or donors/group) was performed on the Fluidigm Biomark HD at the Penn Genomic Analysis Core. A Fluidigm Dynamic Array IFC (BMK-M-96.96) was loaded with pre-amplified cDNA for the 48 samples in duplicate and 20X TaqMan probes for the 96 genes (**Supplementary Table S4**). The probe for TBXT was custom made using the Custom Assay Design tool by ThermoFisher. A brightfield image of the loaded Fluidigm chip was assessed to detect bubbles in the reaction wells. If a bubble was detected, the reaction well was excluded from analysis. To account for variations in total RNA between samples, ΔC_T_ for a sample was calculated by subtracting the duplicate C_T_ average for a gene of interest from the housekeeping gene average (TBP, 18S, and RPS17) for that sample.

To determine the largest effects caused by TP14 treatment, ΔC_T_ values for SCL, +TP, and AF groups were compared. Specifically, genes were plotted for group comparisons if the following conditions were met: (1) the absolute value of the SCL vs AF ΔΔC_T_ was >2, (2) the absolute value of the difference between SCL vs AF ΔΔC_T_ and TP14 vs AF ΔΔC_T_ was >2, (3) there were at least 2 measured ΔC_T_ values for each sample type for each gene of interest (**Fig. 6, Supplementary Figure S7**).

#### Principal Component Analysis (PCA)

PCA and hierarchical clustering were performed using ClustVis^43^. ΔC_T_ for all genes except housekeeping genes were entered into ClustVis. For PCA of SCL and AF samples to identify signature genes for each cell type, only data from these groups were used in the PCA. Plots were generated using custom R scripts with the “ggplot2”, “rgl”, and “Rcmdr” packages.

### Illustrations

Illustrations were generated using Inkscape or BioRender.com.

### Statistical Analysis

The Shapiro-Wilk normality test was used to determine the need for non-parametric testing (alpha = 0.05). For gene expression comparisons at the stages of differentiation leading up to SCL, gene expression was compared via a one-way ANOVA with Tukey’s multiple comparisons post-hoc analysis. For gene expression, nuclear aspect ratio, and SOX9 nuclear intensity comparisons between iPSC and SCL groups, an unpaired t-test with Welch’s correction was used. Gene expression between groups treated with factors for AF-like differentiation of SCL cells was compared to Ctrl groups using a one-way ANOVA with Tukey’s multiple comparisons post-hoc analysis. For comparison of principal component scores and gene expression between SCL and AF groups, an unpaired t-test with Welch’s correction was used. To compare principal component scores and gene expression between SCL, AF, and other treatment groups, a one-way ANOVA was used followed by Tukey’s multiple comparisons post-hoc analysis.

## Acknowledgements

This work was supported by the U.S. Department of Veterans Affairs (RR&D I21 RX003447, RR&D I01 RX002274). The contents do not represent the views of the U.S. Department of Veterans Affairs or the U.S. Government.

We would like to thank the Human Pluripotent Stem Cell Core at the Children’s Hospital of Philadelphia for their donation of the iPSC cell lines CHOP WT4.3 and CHOP WT10.2. We specifically would like to thank Dr. Deborah French and Chintan Jobaliya, M.S. for their knowledge, support, and guidance regarding iPSC culture. In addition, we would like to thank Dr. Farshid Guilak, Dr. Amanda Dicks, and Dr. Chia-Lung Wu from the University of Washington in Saint Louis for their guidance regarding the sclerotome differentiation protocol published by Adkar and colleagues. Finally, we would like to thank the UPenn Genomic Analysis Core for running the Fluidigm GE 96.96 Dynamic Array.

## Author contributions

A.P.P and R.L.M. conceived the experiments. A.P.P. conducted the experiments, performed statistical analysis, and generated figures, schematics, and illustrations. E.B. conducted image analysis using CellProfiler. A.P.P., T.K.T., X.J., and N.A.D. were involved in the preparation and analysis of the Fluidigm GE 96.96 Dynamic Array. H.E.S., S.E.G., N.A.D. and R.L.M. provided scientific and clinical guidance for the design of experiments. A.P.P. and R.L.M. wrote the manuscript, and all authors reviewed the manuscript and approved the final submission.

## Data availability

The data that support the findings of this study are available from the corresponding author, RLM, upon reasonable request.

## Additional Information

None of the authors have conflicts to disclose.

