## Supplementary Information for "Developmental Morphogens Direct Human Induced Pluripotent Stem Cells Towards an Annulus Fibrosus-Like Cell Phenotype"

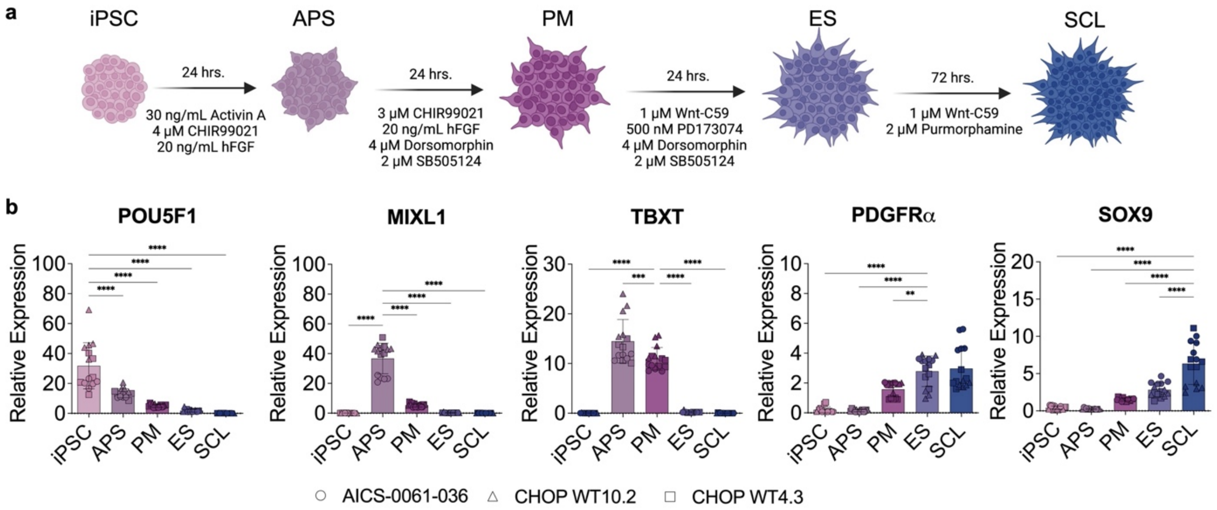


**Supplementary Figure S1. Stepwise differentiation of iPSCs towards the sclerotome fate.** (a) Differentiation protocol used to drive iPSCs towards the SCL fate. (b) Gene expression for the stage-specific genes POU5F1 (iPSC), MIXL1(APS), TBXT (APS and PM), PDGRα (ES and SCL), and SOX9 (SCL), relative to the housekeeping gene, at each stage of differentiation for three iPSC cell lines (n=4/cell line; mean ± s.d.). Statistical significance denoted by * p-value < 0.05, ** < 0.01, *** < 0.001, **** < 0.0001.

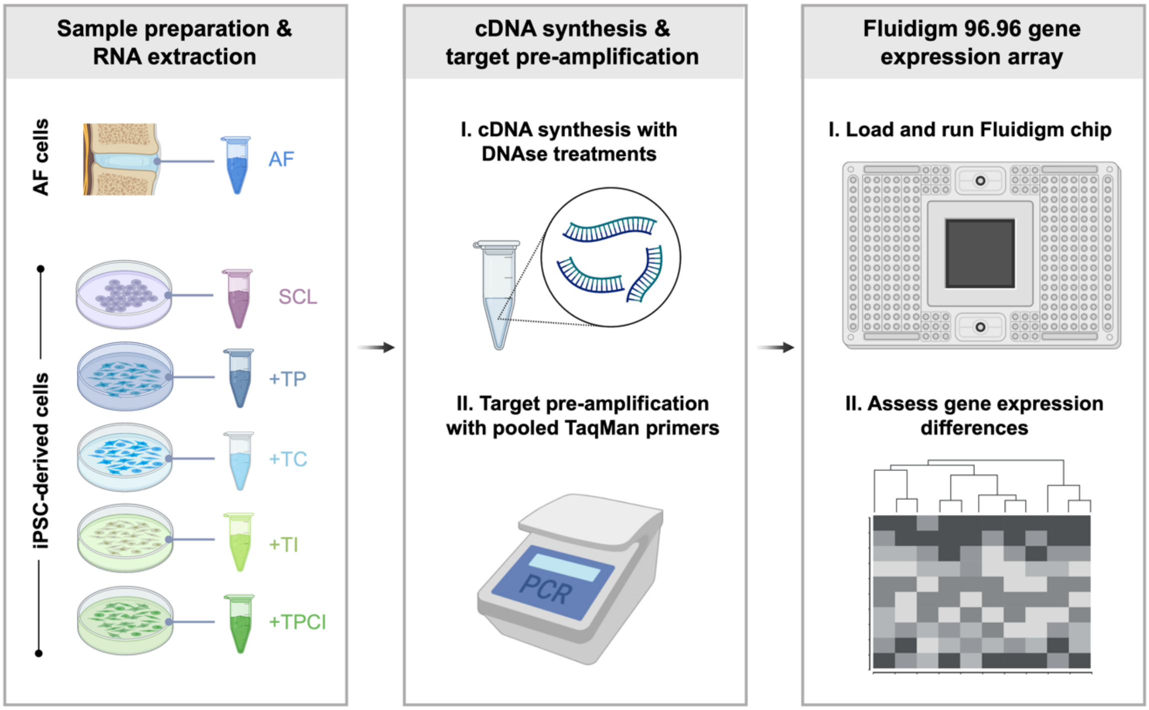


**Supplementary Figure S2. Sample preparation for Fluidigm 96.96 gene expression array.** RNA was extracted from cell samples, after which cDNA was synthesized, and targets were pre-amplified using pooled TaqMan probes. A Fluidigm 96.96 gene expression array was run and the expression of the 96 genes of interest was compared between samples.

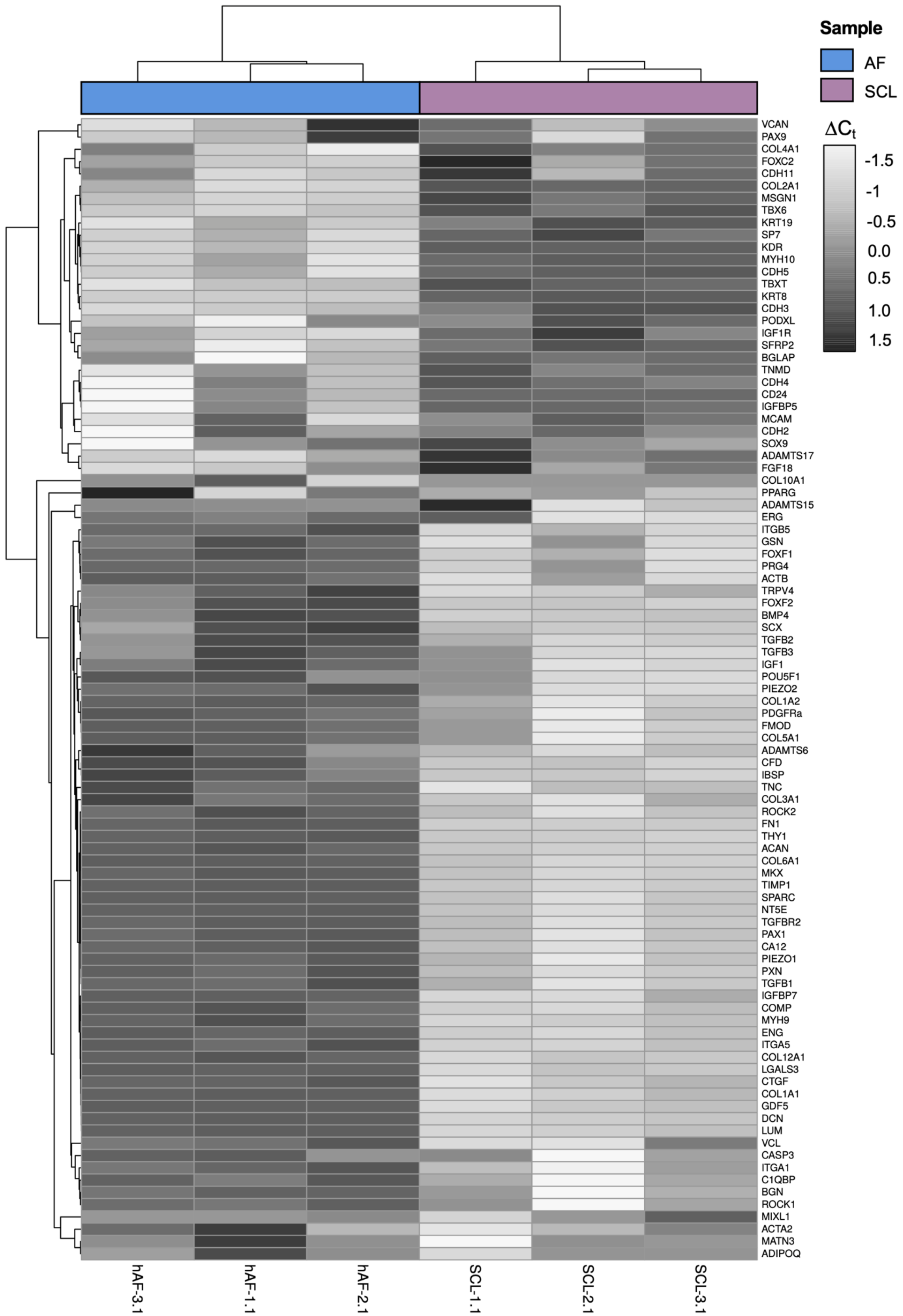
Sample
AF
SCL
ΔCt
-1.5
-1
-0.5
0.0
0.5
1.0
1.5
VCAN
PAX9
COL4A1
FOXC2
CDH11
COL2A1
MSGN1
TBX6
KRT19
SP7
KDR
MYH10
CDH5
TBXT
KRT8
CDH3
PODXL
IGF1R
SFRP2
BGLAP
TNMD
CDH4
CD24
IGFBP5
MCAM
CDH2
SOX9
ADAMTS17
FGF18
COL10A1
PPARG
ADAMTS15
ERG
ITGB5
GSN
FOXF1
PRG4
ACTB
TRPV4
FOXF2
BMP4
SCX
TGFB2
TGFB3
IGF1
POU5F1
PIEZO2
COL1A2
PDGFRa
FMOD
COL5A1
ADAMTS6
CFD
IBSP
TNC
COL3A1
ROCK2
FN1
THY1
ACAN
COL6A1
MKX
TIMP1
SPARC
NT5E
TGFBR2
PAX1
CA12
PIEZO1
PXN
TGFB1
IGFBP7
COMP
MYH9
ENG
ITGA5
COL12A1
LGALS3
CTGF
COL1A1
GDF5
DCN
LUM
VCL
CASP3
ITGA1
C1QBP
BGN
ROCK1
MIXL1
ACTA2
MATN3
ADIPOQ
hAF-3.1
hAF-1.1
hAF-2.1
SCL-1.1
SCL-2.1
SCL-3.1

**Supplementary Figure S3. Heatmap of $\Delta C_t$ values for AF and SCL from Fluidigm gene expression array for 93 genes of interest.** Heatmap of $\Delta C_t$ values for AF vs. SCL PCA, with hierarchical clustering based on the 93 genes included in the gene expression array with color coding by sample type.

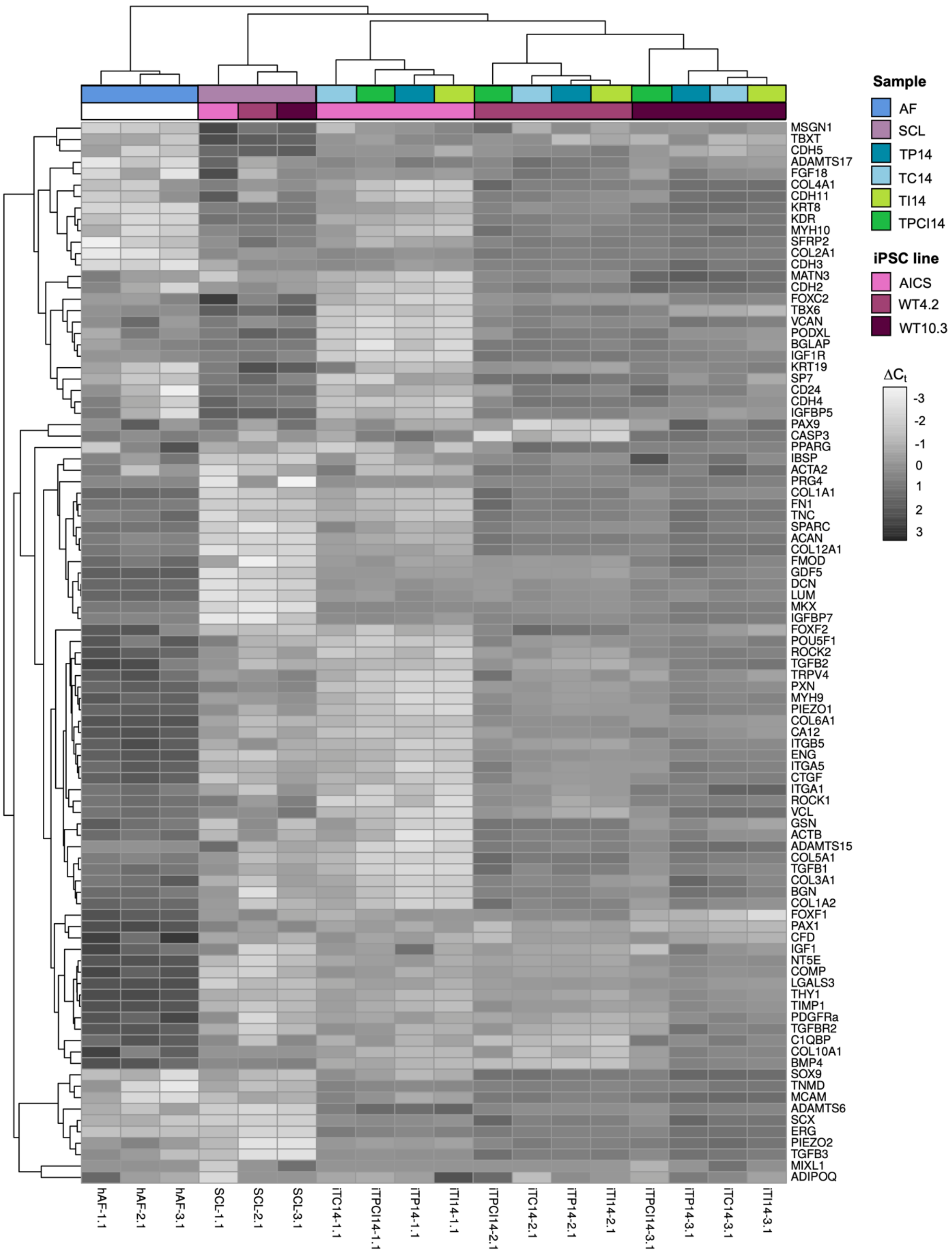


**Supplementary Figure S4. Heatmap of $\Delta C_t$ values for AF, SCL, and all treatment groups from Fluidigm gene expression array for 93 genes of interest.** Heatmap of $\Delta C_t$ values for AF, SCL, and treatment group PCA, with hierarchical clustering based on the 93 genes included in the gene expression array with color coding by sample type and iPSC cell line, where used.

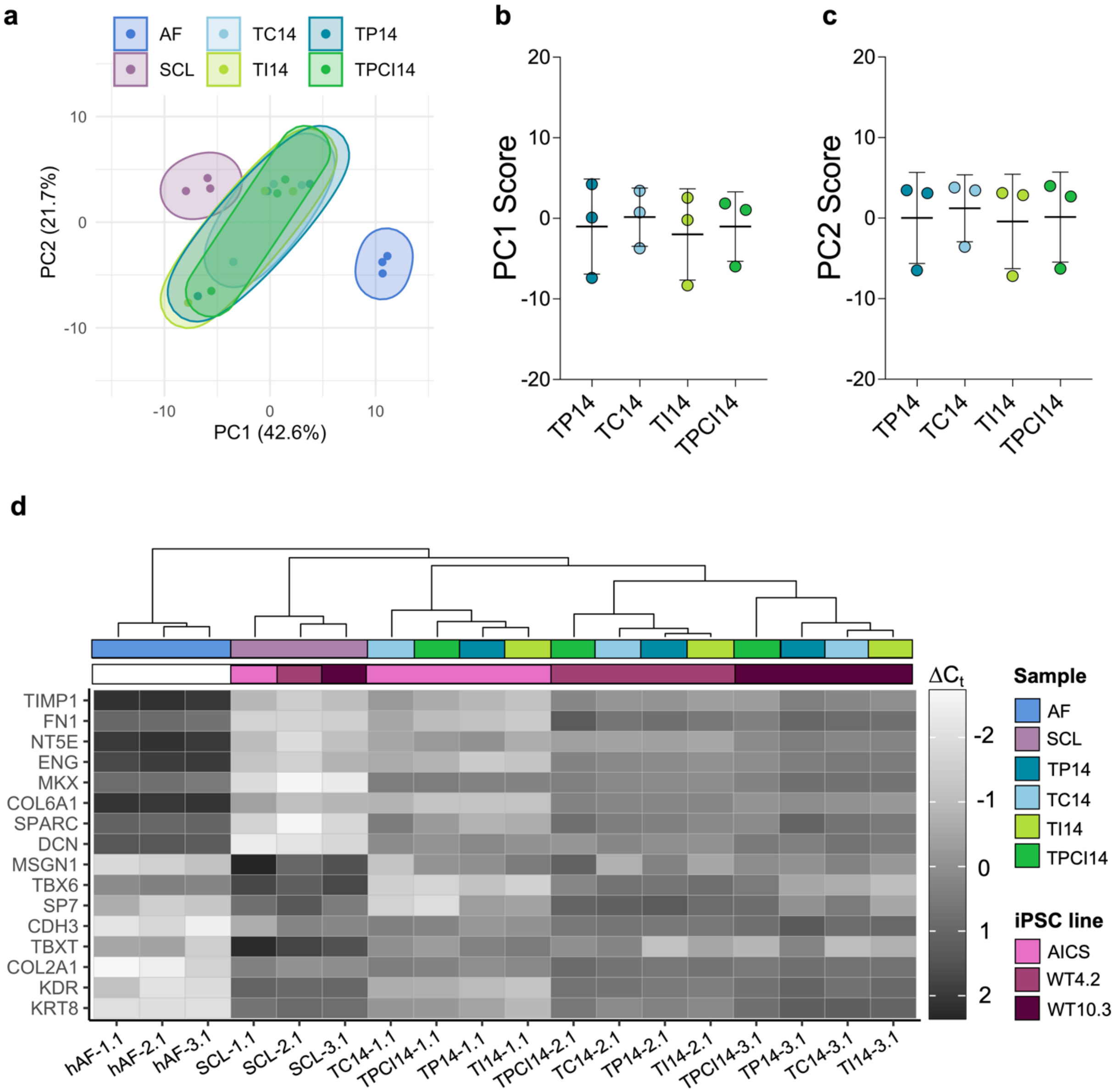


**Supplementary Figure S5. Treatment of SCL cells with factors involved in AF embryonic development increase the expression of genes highly expressed by adult AF cells.** (a) PCA of the data represented by mapping PC1 and PC2 scores for SCL, AF, and factor treatment groups (Ctrl, TP14, TC14, TI14, and TPCI14). Percent of described total variance by each PC is listed in parentheses. Concentration ellipses for each group are demarcated via clustering. (b) PC1 scores and (c) PC2 scores for factor treatment groups (n=3/group; mean ± s.d.). (d) Heatmap of $\Delta C_t$ values for the 8 highest and 8 lowest valued genes identified from the AF vs. SCL PCA, with hierarchical clustering based on the 93 genes included in the gene expression array with color coding by sample type and iPSC cell line. For the heatmap showing all 93 genes used in the cluster analysis, see Supplementary Figure S4. Statistical significance denoted by * p-value < 0.05, ** < 0.01, *** < 0.001, **** <0.0001.

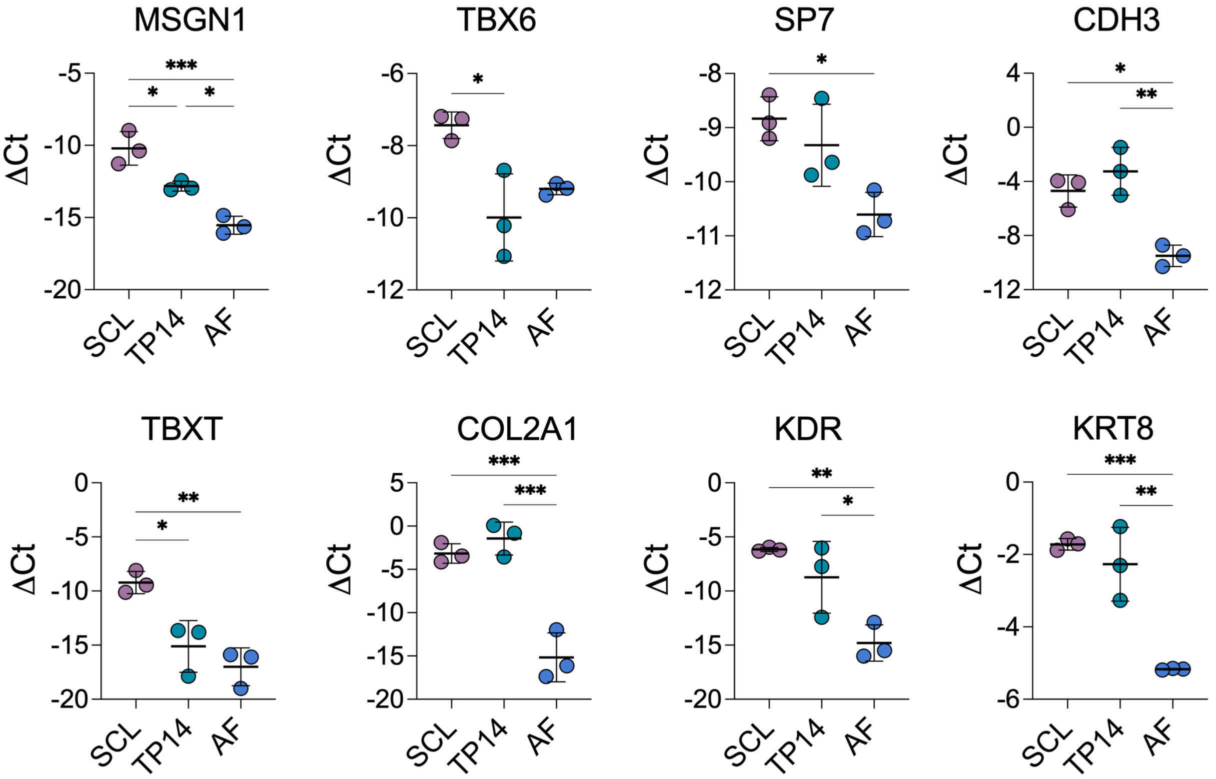


**Supplementary Figure S6. Changes in gene expression driven by factor treatment for genes highly expressed by SCL cells.** $\Delta C_t$ values for genes with lowest PC1 score in AF vs. SCL PCA (highest expression shown by SCL cells) (n=3/group; mean ± s.d.). Statistical significance denoted by * p-value < 0.05, ** < 0.01, *** < 0.001, **** <0.0001.

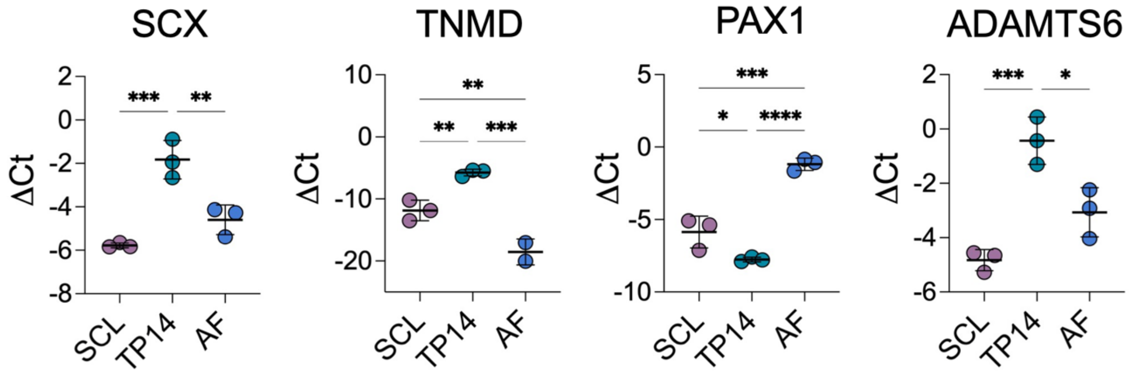


**Supplementary Figure S7. TP14 treatment generates changes in gene expression that are different from mature AF cells.** $\Delta C_t$ values for genes differentially expressed by TP14 cells in comparison to SCL and AF cells (n=3/group; mean ± s.d.). Statistical significance denoted by * p-value < 0.05, ** < 0.01, *** < 0.001, **** <0.0001.

**Supplementary Table S1.** Factors used for differentiation protocols.

| Molecule or GF | Company, Catalog Number | Final Concentration | Resuspended in |
|---|---|---|---|
| SB 505124 | Tocris, 3263 | 2 μM | DMSO |
| PD 173074 | Tocris, 3044 | 500 nM | DMSO |
| Dorsomorphin | Stemgent, 04-0024 | 4 μM | H20 |
| Wnt-C59 | Cellagen Tech, C7641-2s | 1 μM | DMSO |
| Purmorphamine | Stemgent, 04-0009 | 2 μM | DMSO |
| Activin A | R&D, 338-AC-01M | 30 ng/mL | 4 mM HCl + 0.1 %BSA |
| CHIR99021 | Tocris, 4423 | 3-4 mM | H20 |
| bFGF-2 | R&D, 233-FB-01M | 20 ng/mL | PBS + 0.1 %BSA |
| CTGF | Fisher, RD172035100 | 100 ng/mL | Acetate Buffer (0.1 M, pH 4.0) |
| PDGF-BB | Fisher, 220-BB-010 | 2 ng/mL | 4 mM HCl |
| TGF-β3 | R&D, 243-B3/CF | 10 ng/mL | 4 mM HCl + 0.1 % BSA |
| IGF-1 | R&D, 291-G1-200 | 5 ng/mL | PBS |

**Supplementary Table S2.** Custom designed primer sequences for quantitative RT-PCR.

| Target | Forward Primer | Reverse Primer |
|---|---|---|
| POU5F1 | GCAGGAGTCGGGGTGGAGAGCAACT | TCCAGCTTCACGGCACCAGGGGT |
| MIXL1 | CTGAGGAGCCATGACTGACA | TGGGAGTGTGGGCTTAAAAC |
| TBXT | CCCGAAAGATGCAGTGACTT | TACTGCAGGTGTGAGCAAGG |
| PDGFRα | TTCCTGTAACTGGCGGATTC | GCAAGAGGCAACACTGACAA |
| SOX9 | TGACCAGTACCTGCCGCCCAACGG | TGCTGATGCCGTAGCTGCCCGTGTA |
| COL1A1 | CCTACCACTGCAAGAACAGCGTGGCCT | CGATCTCGTTGGAGCCCTGGAGGAGCA |
| COL2A1 | GGGGCAAGACTGTTATCGAG | TGCAACGGATTGTGTTGTTT |
| ACAN | ACAGCTGGGGACATTAGTGG | GTGGAATGCAGAGGTGGTTT |
| ELN | GCATTCCTACTTACGGGGTTG | CTCCGACACCAGGGACAC |

**Supplementary Table S3.** Antibodies and products used for immunofluorescence staining.

| Target | Product | Company, Catalog Number | Dilution |
|---|---|---|---|
| Nucleus | DAPI | ThermoFisher, D1306 | 1:1000 |
| F-actin | Alexa Fluor™ 488 Phalloidin | ThermoFisher, A12379 | 1:500 |
| Sox-9 | SOX9 (D8G8H) Rabbit mAb | Cell Signaling Technology, 82630 | 1:100 |
| Ssea-4 | SSEA-4 Mouse mAb, Clone MC-813-70 | Stemcell Technologies, 60062 | 1:100 |
| E-cadherin | E-cadherin (24E10) Rabbit mAb | Cell Signaling Technology, 3195S | 1:100 |
| Collagen type I | Anti-Collagen I Mouse mAb | Abcam, ab6308 | 1:100 |
| Elastin | Anti-Elastin Rabbit pAb | Abcam, ab21610 | 1:100 |
| Rabbit IgG | Goat anti-Rabbit IgG Alexa Fluor 488 | ThermoFisher, A32731 | 1:1000 |
| Rabbit IgG | Donkey anti-rabbit IgG Alexa Fluor 555 | ThermoFisher, A32794 | 1:1000 |
| Rabbit IgG | Donkey anti-rabbit IgG Alexa Fluor 647 | ThermoFisher, A32795 | 1:1000 |
| Mouse IgG | Goat anti-mouse IgG Alexa Fluor 555 | ThermoFisher, A32723 | 1:1000 |

**Supplementary Table S4.** TaqMan probes used for Fluidigm 96.96 gene expression array.

| Target | Probe |
|---|---|
| TBP | Hs00427620_m1 |
| 18S | Hs99999901_s1 |
| RPS17 | Hs00734303_g1 |
| COL1A1 | Hs00164004_m1 |
| COL1A2 | Hs01028956_m1 |
| COL2A1 | Hs00264051_m1 |
| COL3A1 | Hs00943809_m1 |
| COL4A1 | Hs00266237_m1 |
| COL5A1 | Hs00609088_m1 |
| COL6A1 | Hs01095585_m1 |
| COL10A1 | Hs00166657_m1 |
| COL12A1 | Hs00189184_m1 |
| ACAN | Hs00153936_m1 |
| BGN | Hs00959143_m1 |
| DCN | Hs00370384_m1 |
| FMOD | Hs00157619_m1 |
| TNMD | Hs00223332_m1 |
| LUM | Hs00929860_m1 |
| VCAN | Hs00171642_m1 |
| KRT19 | Hs00761767_s1 |
| PODXL | Hs01574644_m1 |
| POU5F1 | Hs04195369_s1 |
| MIXL1 | Hs04400364_m1 |
| KDR | Hs00911700_m1 |
| TBXT | Custom Ordered |
| MSGN1 | Hs03405514_s1 |
| FOXC2 | Hs00270951_s1 |
| PDGFRa | Hs00998018_m1 |
| PAX1 | Hs01071293_g1 |
| PAX9 | Hs00196354_m1 |
| SCX | Hs03054634_g1 |
| SOX9 | Hs00165814_m1 |
| TBX6 | Hs00365539_m1 |
| MKX | Hs00543190_m1 |
| ADAMTS17 | Hs01020505_m1 |
| SFRP2 | Hs00293258_m1 |

| | |
|---|---|
| FN1 | Hs01549976_m1 |
| ADAMTS15 | Hs00373520_m1 |
| ADAMTS6 | Hs01058097_m1 |
| FOXF1 | Hs00230962_m1 |
| FOXF2 | Hs00230963_m1 |
| CASP3 | Hs00234387_m1 |
| MCAM | Hs00174838_m1 |
| ACTA2 | Hs00426835_g1 |
| ROCK1 | Hs01127699_m1 |
| ROCK2 | Hs00178154_m1 |
| MYH9 | Hs00159522_m1 |
| MYH10 | Hs00992055_m1 |
| GSN | Hs00609272_m1 |
| ACTB | Hs99999903_m1 |
| VCL | Hs00419715_m1 |
| PXN | Hs01104424_m1 |
| ITGA5 | Hs01547673_m1 |
| ITGA1 | Hs00235006_m1 |
| ITGB5 | Hs00174435_m1 |
| PIEZO1 | Hs00207230_m1 |
| PIEZO2 | Hs00926218_m1 |
| TRPV4 | Hs01099348_m1 |
| CDH2 | Hs00983056_m1 |
| CDH11 | Hs00156438_m1 |
| CDH3 | Hs00999915_m1 |
| CDH4 | Hs00899698_m1 |
| CDH5 | Hs00901463_m1 |
| TGFB1 | Hs00998133_m1 |
| TGFB2 | Hs00234244_m1 |
| TGFB3 | Hs01086000_m1 |
| TGFBR2 | Hs00234253_m1 |
| BMP4 | Hs00370078_m1 |
| GDF5 | Hs00167060_m1 |
| CTGF | Hs00170014_m1 |
| IGFBP7 | Hs00266026_m1 |
| IGF1 | Hs01547656_m1 |
| IGF1R | Hs00609566_m1 |
| KRT8 | Hs01595539_g1 |
| CA12 | Hs01080902_m1 |

| | |
|---|---|
| LGALS3 | Hs00173587_m1 |
| CD24 | Hs02379687_s1 |
| COMP | Hs00164359_m1 |
| C1QBP | Hs00241825_m1 |
| MATN3 | Hs00159081_m1 |
| PRG4 | Hs00981633_m1 |
| TIMP1 | Hs99999139_m1 |
| ERG | Hs01554629_m1 |
| IGFBP5 | Hs00181213_m1 |
| FGF18 | Hs00818572_m1 |
| SPARC | Hs00234160_m1 |
| ADIPOQ | Hs00605917_m1 |
| PPARG | Hs01115513_m1 |
| CFD | Hs00157263_m1 |
| SP7 | Hs01866874_s1 |
| BGLAP | Hs01587814_g1 |
| IBSP | Hs00173720_m1 |
| NT5E | Hs01573922_m1 |
| THY1 | Hs00174816_m1 |
| ENG | Hs00923996_m1 |
| TNC | Hs01115665_m1 |

**Supplementary Table S5.** Ranked PC1 loading values for genes included in the AF vs. SCL PCA.

| Gene | PC1 Loading Value |
|---|---|
| KRT8 | -0.1212177 |
| KDR | -0.1175906 |
| COL2A1 | -0.1158053 |
| TBXT | -0.1155829 |
| CDH3 | -0.115438 |
| SP7 | -0.1154355 |
| TBX6 | -0.11482 |
| MSGN1 | -0.1148073 |
| CDH5 | -0.1136989 |
| MYH10 | -0.1121694 |
| KRT19 | -0.1106561 |
| SFRP2 | -0.110621 |
| IGF1R | -0.1080365 |
| ADAMTS17 | -0.101585 |
| IGFBP5 | -0.0942144 |
| BGLAP | -0.0941441 |
| PODXL | -0.0938976 |
| CD24 | -0.09351 |
| COL4A1 | -0.0897712 |
| TNMD | -0.0862157 |
| CDH4 | -0.0859409 |
| FOXC2 | -0.0842064 |
| FGF18 | -0.078512 |
| MCAM | -0.0715894 |
| CDH11 | -0.0695328 |
| CDH2 | -0.052352 |
| SOX9 | -0.0494553 |
| VCAN | -0.0041455 |
| COL10A1 | 0.00063991 |
| MIXL1 | 0.00883256 |
| PAX9 | 0.00923666 |
| ADAMTS15 | 0.02033219 |
| PPARG | 0.04795372 |
| ADIPOQ | 0.05167883 |
| MATN3 | 0.06958652 |
| ACTA2 | 0.07277765 |

| | |
|---|---|
| ERG | 0.07385976 |
| CASP3 | 0.08768762 |
| PRG4 | 0.09215125 |
| VCL | 0.09637539 |
| ROCK1 | 0.09638603 |
| TGFB3 | 0.09853479 |
| POU5F1 | 0.09930633 |
| GSN | 0.0994513 |
| SCX | 0.10107063 |
| ADAMTS6 | 0.10367649 |
| BGN | 0.10421614 |
| IGF1 | 0.10518346 |
| C1QBP | 0.10543632 |
| ACTB | 0.10638252 |
| TGFB2 | 0.10750999 |
| PIEZO2 | 0.10763445 |
| ITGA1 | 0.10896738 |
| TRPV4 | 0.10948782 |
| BMP4 | 0.11003537 |
| COL5A1 | 0.11007259 |
| FMOD | 0.11059481 |
| TNC | 0.11107455 |
| PDGFRa | 0.1114986 |
| FOXF2 | 0.11157535 |
| CFD | 0.11177936 |
| FOXF1 | 0.11259437 |
| COL3A1 | 0.11311522 |
| IBSP | 0.11347423 |
| ITGB5 | 0.11350506 |
| TGFB1 | 0.11572191 |
| CTGF | 0.11627482 |
| COL1A2 | 0.11664488 |
| PIEZO1 | 0.11712166 |
| IGFBP7 | 0.11745759 |
| ROCK2 | 0.11794844 |
| COL1A1 | 0.11857807 |
| MYH9 | 0.11861391 |
| LGALS3 | 0.11861796 |
| PXN | 0.1186919 |

| | |
|---|---|
| PAX1 | 0.11878448 |
| GDF5 | 0.11887985 |
| COL12A1 | 0.11888097 |
| ITGA5 | 0.11919937 |
| COMP | 0.11944478 |
| TGFBR2 | 0.11967663 |
| CA12 | 0.11980191 |
| LUM | 0.12003349 |
| ACAN | 0.12045461 |
| THY1 | 0.12052615 |
| DCN | 0.12063277 |
| SPARC | 0.12067306 |
| COL6A1 | 0.12072834 |
| MKX | 0.12078269 |
| ENG | 0.12079044 |
| NT5E | 0.12079975 |
| FN1 | 0.12101928 |
| TIMP1 | 0.12130947 |
